# On a non-trivial application of Algebraic Topology to Molecular Biology

**DOI:** 10.1101/240499

**Authors:** Ido Braslavsky, Joel Stavans

## Abstract

Brouwer’s fixed point theorem, a fundamental theorem in algebraic topology proved more than a hundred years ago, states that given any continuous map from a closed, simply connected set into itself, there is a point that is mapped unto itself. Here we point out the connection between a one-dimensional application of Brouwer’s fixed point theorem and a mechanism proposed to explain how extension of single-stranded DNA substrates by recombinases of the RecA superfamily facilitates significantly the search for homologous sequences on long chromosomes.

---

Homologous recombination is a highly conserved molecular process that drives the generation of genetic diversity in both prokaryotic and eukaryotic organisms, and plays a pivotal role in the repair of DNA lesions. For example, during the horizontal gene transfer processes of conjugation and transformation in bacteria, exogenous pieces of DNA thousands bases-long are imported into a bacterial cell and incorporated into the chromosome at specific loci of sufficiently high homology. The search for these loci, as well as the strand exchange process that ensues after homology is found are catalyzed by the RecA protein. Given that bacterial genomes are very long molecules, the number of sequences that must be potentially interrogated until homology is found can be exceedingly large. Yet, the search for homology is completed rapidly (1), leading to the notion of facilitated search processes. The search for homology remains as one of the main puzzles behind homologous recombination processes and continues to be a subject of intense experimental and theoretical inquiry (2–7).

The elementary step during the search for homology consists of the juxtaposition of a RecA-single-stranded DNA nucleoprotein filament in which the single-stranded DNA (ssDNA) is stretched by ^~^50% (8), and B-form double-stranded DNA (dsDNA), forming a triple complex or synapse. It is within a synapse that the two sequences are compared, with the first and rapid homology test involving ~8 bases (2, 9, 10). If the test is successful, strand exchange ensues. Otherwise, as is the case of most events, given the large excess of heterologous DNA, new rounds of homology testing follow.

Different aspects of the molecular encounter between a RecA-ssDNA complex and a dsDNA segment and their mutual interaction, all of which may facilitate in some way the search for a homologous target sequence, have been the subject of theoretical modeling. These include inter-segmental transfer (11), the polymeric nature of the dsDNA chromosome (12), sliding (13), rotation (5), the extension of a ssDNA substrate upon polymerization of RecA (14), and the extension of the dsDNA segment upon interaction with the RecA-ssDNA complex (15).

Here we revisit the effects of extension of a ssDNA substrate by the polymerization of RecA (^~^50%) in facilitating the search for homology on long genomes, as elucidated a little more than ten years ago by Bruinsma and co-workers (14). In a nutshell, their mechanism, illustrated schematically in Fig. 1A, ensures that when a RecA-ssDNA complex and dsDNA are aligned, some position along the molecules will be in local registry. Once a local region of registry is found, nucleation of a strand-exchange reaction can ensue. In contrast, when the ssDNA is not stretched (Fig. 1B), any relative displacement between the two molecules results in total loss of registry. The substantial increase in the persistence length of ssDNA due to binding of RecA precludes initiation of strand exchange at other positions, thereby preventing topological traps (14).

The purpose of the present comment is to point out that the efficient and elegant mechanism of target location proposed by Bruinsma and co-workers is in fact a one-dimensional manifestation of a classic, foundational theorem in algebraic topology, the Brouwer fixed point theorem, proved independently by two mathematicians, Brouwer (16) and Bohl (17) about a hundred years before the work of Bruinsma and colleagues, while an equivalent result was proved even earlier by Poincaré in his studies of the three-body problem and celestial mechanics (18). One statement of the theorem is:

**Brouwer’s Fixed Point Theorem**. Given any continuous function *f: X → X*, where *X* is a convex, compact set in a topological space, then there exists a fixed point, that is, a point *x*_0_ ϵ *X* such that *f*(*x*_0_) = *x*_0_.

Here “convex, compact set” refers to a bounded set that includes its boundary and has no holes. In two dimensions, this theorem ensures that a point in a map of say, a city, lies exactly over its corresponding point when read over the city limits, no matter what the map scale is, or if it is rotated or crumpled. In three dimensions, there will be one fluid particle that returns to its starting point, after stirring the coffee in a cup. In the context of homologous recombination, a one-dimensional case, the (continuous) function is furnished by the stretching of a segment *x* of ssDNA by RecA and its shift, i.e. *f*(*x*) *= kx − shift*, with *k* >1. The existence of the fixed point translates into the local coincidence between one or few bases in the ssDNA and the dsDNA segment, which occurs as long as (*k – 1*)*x_Max_ ≥ shift ≥* 0 (Fig. 1C). The fact that the stretching of the ssDNA by RecA binding is actually not uniform, with bases organized into near B-form triplets (19), does not affect the applicability of Brouwer’s fixed point theorem, as the stretching function is still continuous. The only modification required is that shifts should take place in units of triplets instead of individual bases (Fig. 1D). Non-uniform stretching does not alter the improvement in search time either, which is of order *~N/*((*k –* 1)*L +* l), where *N* is the number of base pairs of the genomic dsDNA, and *L* the number of homologous bases on ssDNA covered by RecA. This of course assumes that both the RecA-ssDNA complex and the dsDNA are straight. In actuality, on large scales genomic dsDNA is highly convoluted as the genome must be highly compacted to fit into a cell (10, 20, 21). The highly convoluted configurations of the chromosome may allow the parallel search for homology on distal segments (3). It is worthwhile pointing out that the ability to shift relatively the stretched RecA-ssDNA and the dsDNA molecules is what allows short-scale sliding to be effective as a dynamic process facilitating the homology search.

Other recombinases of the RecA superfamily such as Rad51 (22) and Dmc1 (23) in eukaryotes, as well as RadA in archaea (19), also stretch their ssDNA substrates by ^~^50%. It is therefore tempting to suggest that extension of an ssDNA substrate, a hallmark of homologous recombination, may have been preserved throughout evolution precisely to facilitate homology search.

It is nothing but outstanding that in addition to its well-known, numerous applications in various fields of pure Mathematics, Game Theory and Economics, Brouwer’s fixed point theorem has now also found its way into Molecular Biology.

## Acknowledgements.

We thank O. Raz and O. Feinerman for discussions. J.S. is the incumbent of the Siegfried and Irma Ullman Professorial Chair.

---

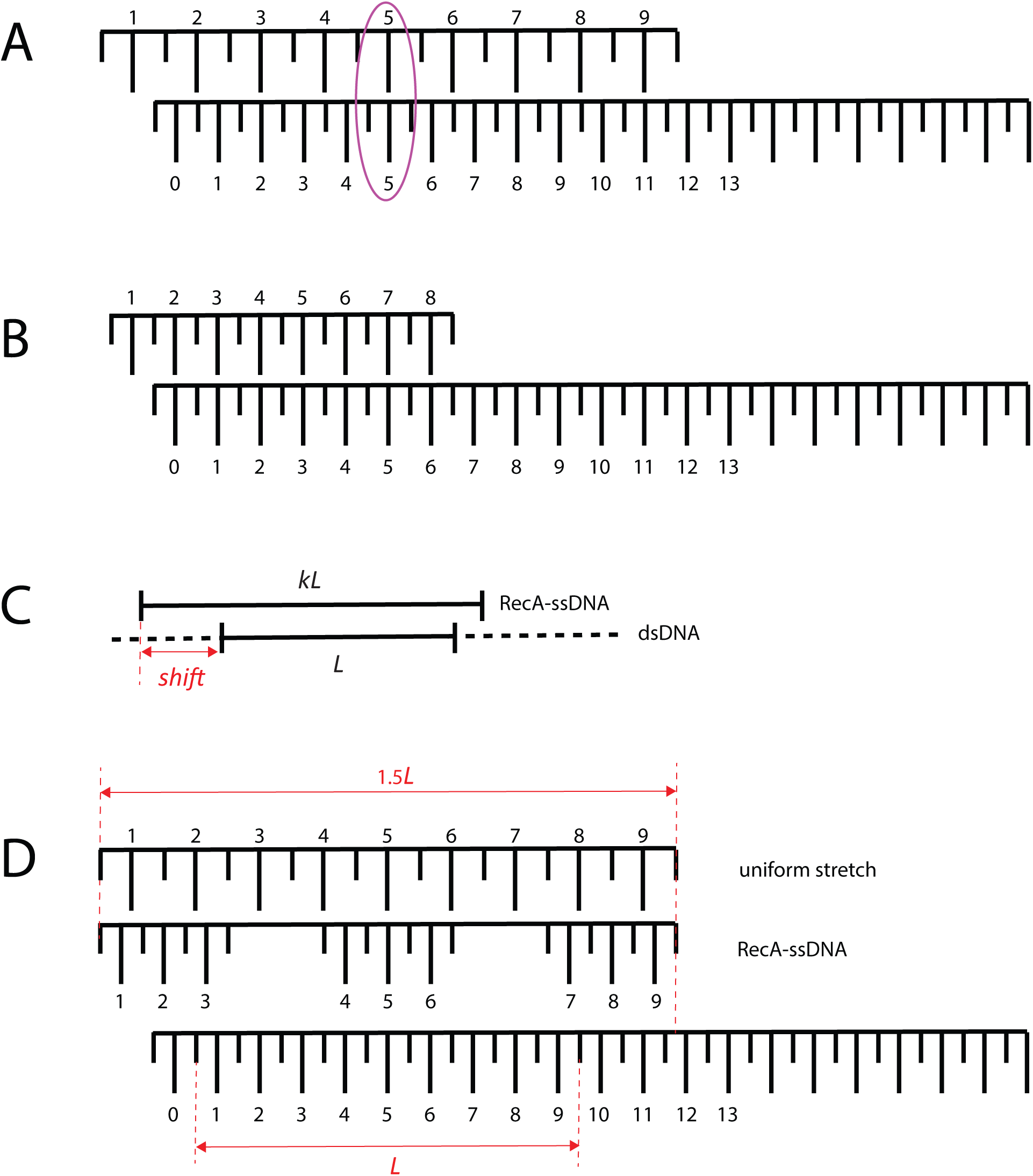
Illustration of Brouwer fixed point theorem in the context of one-dimensional homologous recombination. **(A)** Local homology match between individual bases in a uniformly stretched RecA-ssDNA complex represented by the upper ruler, and a corresponding base on a long, unstretched dsDNA represented by the lower ruler. The ssDNA substrate has been stretched by 50%. (**B**) Arbitrary relative displacements of two rulers both of which are un-stretched, lead to full lack of homology along the entire length of the ssRNA (upper ruler). (**C**) Parameters in the juxtaposition of homologous sequences of ssDNA and dsDNA tracts. (**D**) Local homology match between triplets in a non-uniformly stretched RecA-ssDNA complex in which base triplets preserve their B-form, and a triplet on un-stretched dsDNA.

